# SAP expressing T peripheral helper cells identify systemic lupus erythematosus patients with lupus nephritis

**DOI:** 10.1101/2023.11.16.566902

**Authors:** Yevgeniya Gartshteyn, Laura Geraldino-Pardilla, Leila Khalili, Shoiab Bukhari, Shalom Lerrer, Anca D. Askanase, Adam Mor

**Affiliations:** Division of Rheumatology, Department of Medicine, Columbia University Medical Center, New York, NY, USA; Columbia Center for Translational Immunology, Columbia University Medical Center, New York, NY, USA

**Keywords:** Systemic Lupus, Lupus Nephritis, T peripheral helper cells, T follicular helper cells, SLAM, SAP

## Abstract

**Introduction:** T follicular (TFH) and peripheral helper (TPH) cells have been increasingly recognized as a pathogenic subset of CD4 T cells in systemic lupus erythematosus (SLE). The SLAM Associated Protein (SAP) regulates TFH and TPH function by binding to the co -stimulatory signaling lymphocyte activation molecule family (SLAMF) receptors that mediate T cell – B cell interactions. SAP and SLAMF are critical for TPH dependent B cell maturation into autoantibody-producing plasma cells that characterize SLE pathogenesis.

**Methods:** Peripheral blood mononuclear cells (PBMCs) were isolated using density gradient separation from whole blood. Cells were stained for cell surface markers, followed by permeabilization and staining of intracellular SAP for spectral flow cytometry analysis. Additionally, we analyzed SAP expression from renal infiltrating lupus nephritis (LN) T cells using the publicly available single-cell RNA sequencing (sc-RNA seq) Accelerated Medicines Partnership (AMP) in SLE dataset.

**Results:** PBMCs from 30 patients with SLE (34±10 years old, 83% female), including 10 patients with LN, were analyzed. We found an increase in total SAP-positive CD4 and CD8 T cells in SLE compared with controls (55.5±2.6 vs. 41.3±3.4, p=0.007 and 52.5±3.0 vs. 39.2±2.8, p=0.007 respectively). In CD4 T cells, the highest SAP expression was in the TPH subset. The frequency of SAP+TPH in circulation correlated with disease activity, SLE patients with renal disease had higher levels of circulating SAP+TPH that remained significant after adjusting for age, sex, race, low complements, and elevated anti-dsDNA (p=0.014). scRNA-seq data of renal infiltrating T cells identified increased SAP from LN compared with control kidney biopsy samples (p=0.03), including an expansion of SAP-positive TFH-like subsets in the LN kidneys. Increased SAP expression in LN was associated with the differential expression of SLAMF3 and SLAMF7 as well as granzyme K and EOMES. The existence of two predominant SAP-expressing subsets, the TFH-like CD4 T cells and granzyme K positive effector CD8 T cells, was verified using scRNA-seq data from a human transcriptomic atlas of fifteen major organs.

**Conclusion:** The expansion of SAP expressing T helper cells was associated with lupus nephritis in our cohort and verified using scRNA-seq data of renal infiltrating T cells. Improved understanding of SLAM/SAP signaling can identify new therapeutic targets in LN.

## Introduction

Systemic lupus erythematosus (SLE) is a heterogeneous autoimmune disease with multifactorial pathogenesis and variable organ involvement. B lymphocyte dysregulation and auto-antibody production are essential to the development of SLE. T lymphocytes provide help to autoreactive B cells, supporting their maturation into mature B cells and autoantibody producing plasma cells. Thus, understanding the molecular mechanisms by which CD4 T cells support B cell development is critical for understanding SLE pathogenesis.

The primary site of T-B cell differentiation is in the germinal centers of lymphoid follicles. In SLE, T-B cell interactions also occur in extra-follicular germinal centers that arise within inflamed tissues. T-B cell interactions begin following ligation of the T cell receptor (TCR) by cognate antigen presented by the B cell major histocompatibility complex (MHC). Signaling downstream of TCR-MHC binding is further modulated by secondary activating or inhibitory co-receptors that can enhance or lessen cell activation. The signaling lymphocyte activation molecule family (SLAMF) cell surface receptors, which consists of nine transmembrane proteins (SLAMF1-9), are expressed on both T and B cells and serve as co-stimulatory molecules stabilizing T-B cell interactions in the germinal centers. Genetic loci encoding the SLAMF genes are associated with SLE risk, particularly polymorphisms in SLAMF3, SLAMF4, and SLAMF6.^1–3^ However, only small differences in cell surface SLAMF expression levels have been described in SLE as compared to controls, and these are not consistently seen across studies.^4–6^ We found that SLAMF3 and SLAMF6 expression can be detected on 98% of circulating T cells in SLE and healthy controls.^7^ Therefore, we hypothesized that the receptor’s cell-surface expression does not regulate SLAM signaling and instead SLAM’s role in SLE is modulated by downstream adaptors. In T cells, SLAM receptor signaling is mediated by an adaptor molecule, SLAM Associated Protein (SAP). The binding of SAP to SLAMF receptors in T cells recruits signaling kinases, leading to enhanced activation of TCR and downstream NF-kB pathways.^8, 9^ [The SAP gene (SH2D1a) is located on the X-chromosome, and the absence of SAP results in an X-linked lymphoproliferative disease, XLP, an immunodeficiency syndrome characterized by unstable T-B cell interactions, absence of germinal centers, and lack of isotype switched B cells.^10^ On the other hand, a loss of function SAP frameshift mutation in mouse models of SLE protected the mice from developing lupus, specifically the production of autoantibodies and the development of glomerulonephritis.^11^

T follicular helper cells (TFH), a subset of the CD4 T helper cells, function in the secondary lymphoid organs to promote B cell differentiation. Recognized as PD-1^+^CXCR5^+^CD4^+^ T cells, TFH are recruited to the lymphoid follicles, where they promote B cell maturation through direct T-B cell contact via co-stimulatory receptors and cytokine secretion. T peripheral helper cells (TPH) are a more recently identified subset of CD4 T cells that are transcriptionally and functionally similar to TFH. TPH cells, defined as PD-1^HIGH^CXCR5^-^ CD4^+^, were first isolated from the joints of patients with rheumatoid arthritis and have since been shown to be expanded in lupus and other autoimmune conditions.^12–14^ TPH maintain the ability to support B cell development and antibody production. Still, instead of localizing to the secondary lymphoid tissues, these cells can be isolated from the blood and peripheral tissues, especially at sites of inflammation.

SAP deficiency results in unstable T-B cell interactions and absence of antigen specific antibodies.^15, 16^ In contrast, SLE is a disease of increased ectopic germinal center formations and class switched, long-lived B cells that reflect stable TFH-B cell interactions. This led us to hypothesize that SLAMF6/SAP signaling, especially in the TFH and TPH cells, is enhanced in SLE and contributes to its pathogenesis.

## Material and methods

### Patients

The study population was a sample of 30 adult patients randomly recruited from the Columbia University Lupus Cohort. All patients met the 1997 American College of Rheumatology (ACR) classification criteria. Demographic information was obtained from chart review. Serological covariates were measured at the New York Presbyterian Hospital clinical laboratory and ascertained from chart review; blood specimens for clinical and research use were drawn at the same time. The study was approved by the Columbia University Institutional Review Board.

Eighty three percent were female, mean age was 34±10 (Tab. 1). Ten percent of the patients were Caucasian, 27% were African American and 63% were Hispanic. According to New York City Department of Health neighborhood population estimates (modified from US Census Bureau population estimates), the racial/ethnic distribution of the Manhattan population in 2022 was 31.9% non-Hispanic white, 28.9% Hispanic, 23.4% non-Hispanic black, 14.2% non-Hispanic Asian and 1.6% other. However, the area around Columbia University is known to have a larger percentage of people of Hispanic ethnicity and African descent. For all of these reasons, despite random recruitment of sequential consenting patients, our study population has a greater proportion of minority groups than population data alone would predict.

**Table 1.** Patient Characteristics.

|  | SLE (all patients) |  |  | HC (n=13) |
| --- | --- | --- | --- | --- |
|  | no LN (n=20) | LN (n=10) | p-value |  |
| Age | 35.9 ± 11.4 | 31.2 ± 6.9 | 0.24 | 30.3± 6.2 |
| Sex (Female) | 19 (95%) | 6 (60%) | **0.03** | 10 (78%) |
| Race |  |  |  |  |
| White | 2 (10%) | 1 (10%) | 1.00 | 6 (46%) |
| Black | 6 (30%) | 2 (20%) | 0.68 | 2 (15%) |
| Hispanic | 12 (60%) | 7 (70%) | 0.70 | 2 (15%) |
| Other |  |  |  | 3 (23%) |
| Weight (kg) | 75.0 ± 21.4 | 83.3 ± 35 | 0.71 | -- |
| Ever Smoker | 2 (10%) | 2 (20%) | 0.58 | -- |
| Disease Duration (years) | 3[1-11] | 10.5[4-15] | 0.26 | -- |
| non-renal SLEDAI-2K | 6[3-8] | 5[2-8] | 0.59 | -- |
| ACR Criteria |  |  |  |  |
| malar rash | 11 (55%) | 7 (70%) | 0.69 | -- |
| discoid rash | 6 (30%) | 3 (30%) | 1.00 | -- |
| photosensitivity | 10 (50%) | 2 (20%) | 0.24 | -- |
| mucositis | 8 (40%) | 5 (50%) | 0.71 | -- |
| arthritis | 17 (85%) | 10 (100%) | 0.53 | -- |
| serositis | 9 (45%) | 4 (40%) | 1.00 | -- |
| hematological | 7 (35%) | 6 (60%) | 0.26 | -- |
| Serologies |  |  |  |  |
| ANA | 20 (100%) | 10 (100%) | -- | -- |
| anti-dsDNA | 17 (85%) | 9 (90%) | 0.60 | -- |
| anti-Smith | 7 (35%) | 5 (50%) | 0.46 | -- |
| aPL | 8 (40%) | 3 (33%) | 1.00 | -- |
| Medication |  |  |  |  |
| HCQ | 11 (55%) | 6 (60%) | 0.55 | -- |
| Mycophenolate Mofetil | 4 (20%) | 5 (50%) | 0.11 | -- |
| Calcineurin Inhibitors | 0 | 2 (20%) | 0.10 | -- |
| Biologics (anti-B cell) | 1 (5%) | 1 (1%) | 0.56 | -- |
| Cyclophosphamide | 0 | 1 (1%) | 0.33 | -- |
| Prednisone (>10mg/day) | 5 (25%) | 4 (40%) | 0.33 | -- |

aPL antiphospholipid antibodies; HCQ hydroxychloroquine; HC healthy control

Ten patients had kidney biopsy confirmed lupus nephritis (LN). The median time from kidney biopsy to study enrollment was 228 days (IQR 5-1983 days). Isolated Class IV and Class V was present in 2 and 3 of the patients respectively; the remaining 5 patients had an overlap of class IV/V LN. LN was associated with male sex (p=0.02) and conferred increased disease activity as demonstrated by higher SLEDAI-2K scores.

### General Reagents

RPMI medium 1640, PBS and FBS were purchased from Life Technologies. Staphylococcus Enterotoxin E (SEE) was acquired from Toxin Technology. All stimulations were performed with purified anti-human CD3 (BioLegend #300465), anti-human SLAMF6 (BioLegend #317202) and anti-human SLAMF3 (BioLegend #326109).

### Cell isolation

Peripheral blood mononuclear cells (PBMC) from SLE patients and healthy controls were isolated from peripheral blood using Ficoll gradient centrifugation. Red blood cells were lysed using a lysis buffer (Gibco #A1049201). Primary CD3^+^ T and B cells were isolated using RosetteSep T cell enrichment kit (STEMCELL #15021C) and B cell enrichment kit (STEMCELL #15024C), respectively. All cells were stored in -80C until analysis.

#### Cell stimulation

For immobilized, plate-bound stimulation 48 well plates were coated with anti-CD3 1.5ug/mL and either anti-SLAMF6, anti-SLAMF3 or inactive IgG isotype control at a concentration of 5ug/mL. PBMCs were incubated for 12-18 hours at a density of 500,000 cells per well. For soluble stimulation, primary autologous T and B cells were co-cultured in a 2:1 ratio in the presence of SEE 150pg/mL. anti-SLAMF6 or anti-SLAMF3 antibodies were added to the co-cultures at a concentration of 10ug/mL.

### Spectral flow cytometry analysis

Viability was assessed using Zombie UV dye (BioLegend #423107) and nonspecific Fc interactions were blocked (BioLegend #422302). Cells were stained for cell surface markers as follows: APC/Cy7 anti-CD3 (BioLegend #300318), AF700 anti-CD4 (BioLegend #300526), BV605 anti-CD8a (BioLegend #301040), BV421 anti-PD1 (BioLegend #329920), PE/Cy7 anti-CXCR5 (BioLegend #356924), BV711 anti-CD127 (BioLegend #351328), PE/Cy5 anti-CD25 (BioLegend #302608), PE anti-HLA-DR (BioLegend #307606) and BV510 anti-CD69 (BioLegend #310935). Brilliant Stain Buffer was obtained from BD Biosciences (#563794). Following staining, cells were fixed (BioLegend #420801) and permeabilized (BioLegend #421002) followed by intracellular SAP staining (eBioscience #50-9787-42). Samples were analyzed in five batches containing SLE and healthy control samples. A single reference control sample from the same donor was analyzed in each set to ensure internal validity. Data was acquired on the 5-Laser Cytek Aurora and analyzed using FlowJo v.10.9. Gating strategy is provided in the supplement. (Sup. Fig. 1).

### Sequencing data analysis

We used the single cell RNA sequencing (scRNA-seq) data from SLE kidney biopsy samples, publicly available through the Accelerated Medicines Partnership (AMP) national collaboration (accession number SDY997).^17^ Briefly, leukocytes isolated from kidney biopsies from 24 patients with LN and 10 control samples, acquired from living donor kidney biopsies, were analyzed by scRNA-seq and subjected to stepwise cell clustering to identify cell-specific populations within the kidney. We accessed the scRNA-seq data using the Bioturing Talk2Data platform. Using the original authors’ definitions of the clusters, we compared the expression of the SAP gene, SH2D1A, in T cells from SLE vs. control biopsy samples.

Additionally, we accessed scRNA-seq data sequenced from fifteen major organs of a single healthy volunteer (accession number GSE159929).^18^ Briefly, high-throughput sequencing was performed on viable single cells from the tissue samples of 15 different organs of a research-consented adult donor, ultimately achieving an average of 6,000 cells for each organ. We restricted our analysis to T cells and used normalized gene counts as reported by the Bioturing platform. Dimension reduction was performed using the Uniform Manifold Apporoximation and Projection (UMAP) algorithm. Clustering was performed using the Louvain method.

### Statistical Analysis

Continuous variables were summarized as mean ± standard deviation for normally distributed variables or median ± interquartile range (IQR) for non-normally distributed variables and analyzed using a Student’s t-test or Man-Whitney U respectively. Categorical variables were summarized as frequencies and analyzed using Fischer Exact test. Multivariable modeling was done in Stat/IC 15.1.

Differential gene expression was calculated using the T-test statistic method, with log2FC ≥ 0.6 and a false discovery rate (FDR) ≥ 0.05. Data was plotted using the ggvolcano shiny app. Pathway analyses of enriched genes were assessed using the Kyoto Encyclopedia of Genes and Genomes (KEGG) and Gene Ontology (GO) resources.

## Results

### SAP levels are increased in T Cells from SLE patients

Using spectral flow cytometry, we analyzed the intracellular expression of SAP in circulating, non-stimulated T cells. We found an increase in total SAP-positive CD4 and CD8 T cells in SLE as compared with controls (55.5±2.6 vs. 41.3±3.4, p=0.007 and 52.5±3.0 vs. 39.2±2.8, p=0.007 respectively). Similarly, SAP staining intensity was more significant in CD4 and CD8 T cells in SLE (MFI 6750.8±423.3 vs. 5065.9±488.6, p=0.02 and 5301.8±311.6 vs. 4069.9±265.1, p=0.05 respectively) (Fig. 1A). To better understand whether SAP expression level was associated with an activated T cell phenotype, we looked at the association between SAP levels and HLA-DR and CD69 expression in circulating T cells (Fig. 1B). We found a positive correlation between SAP expression and HLA-DR (Spearman’s rho=0.58, p<0.001) as well as between SAP levels and CD69 (Spearman’s rho=0.40, p=0.01). We therefore identified that a higher percentage of peripheral T cells from SLE patients, compared to healthy controls, express SAP and that these circulating T cells further express markers suggesting recent activation.

**Figure 1.**
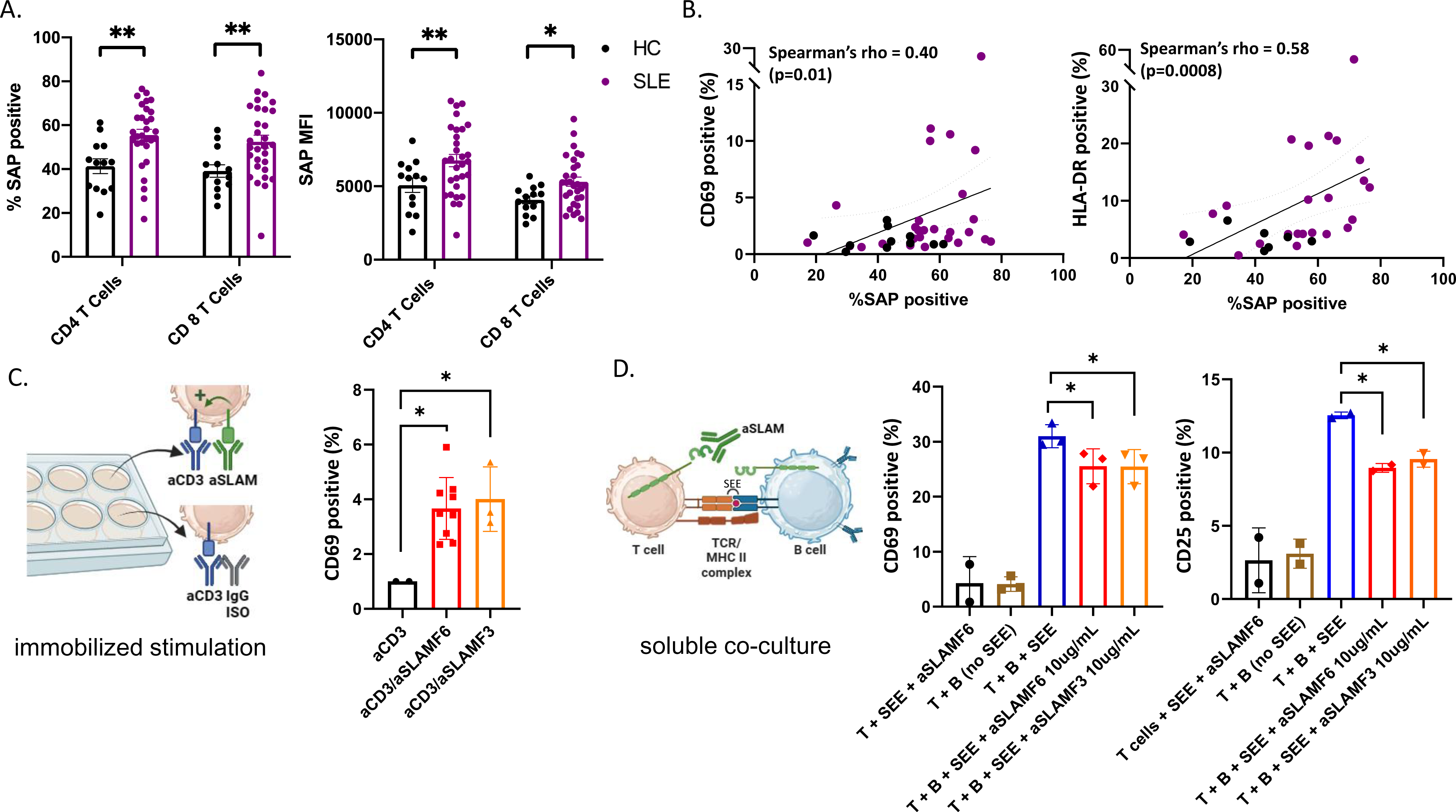
SAP levels are increased in T cells isolated from SLE patients and function to enhance T cell activation. **A)** Peripheral blood mononuclear cells (PBMC) from SLE patients (n=30) and healthy controls (n=13) were evaluated by flow cytometry for SAP expression in T cells, summarized as percent of SAP positive T cells (%SAP positive) or median SAP MFI per person. **B)** Levels of cell surface CD69 (left) and HLA-DR (right) expression on circulating CD4 T cells were quantified and plotted against SAP expression levels, each dot represents a study subject. **C)** Primary T cells were stimulated in-vitro for 4-6 hours with either anti-CD3 (aCD3) alone or aCD3 and anti-SLAMF6 (aSLAMF6) or anti-SLAMF3 (aSLAMF3) (plate-bound stimulation). Following stimulation cells were collected and assessed for CD69 cell surface expression. **D)** Primary autologous T and B cells were co-cultured for 12-16 hours in presence of staphylococcal superantigen (SEE)(soluble co-culture). aSLAMF6 or aSLAMF3 antibodies were added to the co-cocultures to disrupt SLAMF receptor binding in the T-B immunological synapse. Cells were then collected and assessed for CD69 and CD25 cell surface expression. Statistical comparisons were done using a two-sided, unpaired t-test. *p<0.05, **p<0.01, ***p<0.001.

To better understand the functional role of SAP in SLE, we focused on the SLAMF receptor signaling pathways dependent on SAP binding. Given that polymorphisms in SLAMF3 and SLAMF6 are associated with SLE risk, we chose to focus on these two SLAMF receptors for our functional experiments. SAP is a SLAM adaptor protein with two ITSM binding sites, one of which binds to SLAM family receptors, and the other recruits signaling kinases, ultimately coupling SLAM signaling to TCR activation.^9^ We have previously shown that in healthy T cells, signaling through the SLAMF6 receptors enhanced T cell activation.^19^ To confirm SLAMF-mediated T cell activation for another SLAMF receptor and to validate this finding in samples isolated from SLE antibodies, we stimulated peripheral blood mononuclear cells isolated from SLE patients with anti-CD3 antibodies alone or in the presence of activating anti-SLAMF6 or anti-SLAMF3 antibodies (immobilized stimulation, Fig. 1C). We found that T cells activated with anti-CD3 + anti-SLAMF6 or anti-CD3 + anti-SLAMF3 expressed higher levels of CD69 than T cells activated with anti-CD3 alone.

Similarly, we would expect that in circulation homotypic binding of SLAMF receptors expressed by both T and B cells (i.e. SLAMF6 or SLAMF3) supports cell-cell interaction and enhances MHC-TCR activation. We thus asked whether anti-SLAMF6 or anti-SLAMF3 antibodies added to a T-B cell co-culture can disrupt T-B cell SLAMF-SLAMF binding and thus dampen T cell activation (soluble stimulation, Fig. 1D). Indeed, we found that adding either anti-SLAMF6 or anti-SLAMF3 antibodies to an autologous T-B cell co-culture inhibited the expression of CD69 and CD25 on T cells. We thus show that inhibition of SLAMF receptors with competitive monoclonal antibody binding can weaken T-B cell interactions.

### SAP^+^TPH levels are expanded in SLE and are associated with renal involvement

SAP expression in T cells is critical for T cell-mediated B cell maturation.^10, 16^ This led us to evaluate the expression of SAP in CD3^+^CD4^+^PD1^+^CXCR5^+^ TFH cells and the CD3^+^CD4^+^PD1^+^CXCR5^-^ TPH cells. In the SLE donors, TFH cells and TPH cells constituted 2.3±2.8% and 18.7±9.7% of the CD4 T cells. We found that the number of T cells with detectable SAP levels, as well as the median SAP expression level, was significantly greater in TPH population (80%, MFI 10537) as compared to either the PD1^-^CXCR5^-^ T cell subset or the CD127^lo^CD25^hi^ T regulatory cell subset (51%, MFI 6128 and 55%, MFI 6486 respectively, p<0.0001). On the other hand, we found considerable variability across subjects in SAP expression levels in TFH cells, with no significant difference seen in the TFH population as compared to the PD1^-^CXCR5^-^ or the T regulatory cells (Fig. 2A). Thus we found that SAP expression is not uniform across all T cells and that the TPH subset is particularly enriched in SAP expression.

**Figure 2.**
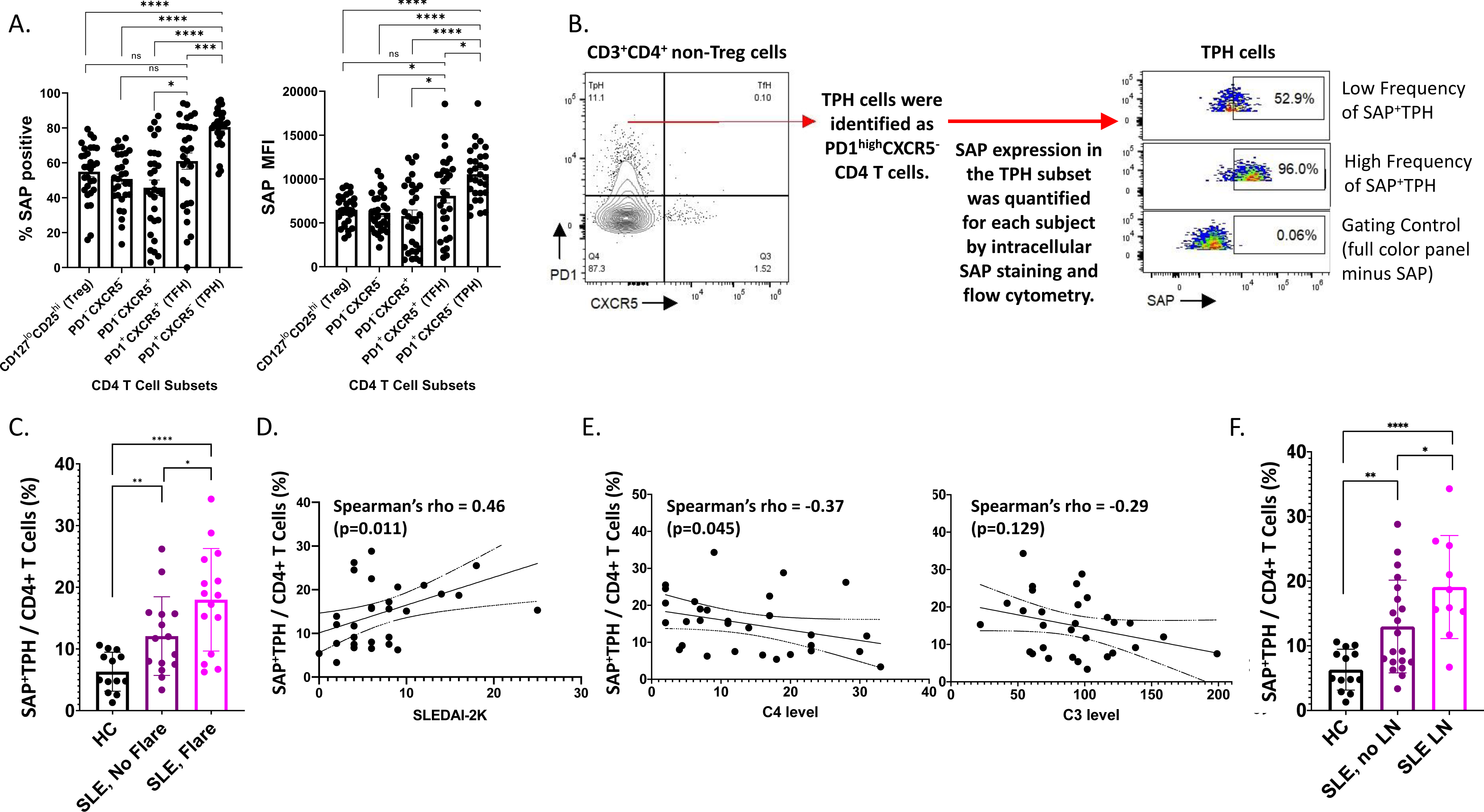
SAP positive TPH cells are expanded in SLE, particularly in biopsy-proven lupus nephritis (LN). **A)** PBMC samples from SLE donors were analyzed by flow cytometry to identify CD4 and CD8 T cell subsets in circulation. In the CD4 T cell subsets, T regulator cells (Treg) were gated out and the remaining CD4 T cells were further sub-divided based on PD1 and CXCR5 expression. The average frequency of SAP expression and median SAP MFI were summarized by CD4 subsets. **B)** TPH cells, defined as PD1^high^CXCR5^-^ were further analyzed based on SAP expression, identifying a new subset of SAP expressing TPH cells (SAP^POS^TPH). **C)** The frequency of SAP^POS^TPH cells in healthy controls, stable SLE, and SLE with disease flare was quantified and compared. **D)** We evaluated for a correlation between individual SAP^POS^TPH levels and SLEDAI-2K clinical disease activity scores and between **E)** SAP^POS^TPH and complement C4 and C3 levels. **F)** The frequency of SAP^POS^TPH cells in healthy controls, non-renal SLE and SLE with LN was quantified and compared. Statistical comparisons were done using a two-sided, unpaired t-test. *p<0.05, **p<0.01, ***p<0.001, ****p<0.0001. The images were created with BioRender.com.

Given the correlation of SAP expression with markers of recent T cell activation, we hypothesized that SAP-positive TPH may identify a subset of TPH cells that more directly contribute to the dysregulated humoral immunity and clinical outcomes in SLE. We defined these cells as SAP-positive TPH or SAP^+^ TPH (Fig. 2B). SAP^+^ TPH cells constituted 6.3±3.0% of all CD4 T cells in the healthy controls and were expanded to 11.8±6.3% in SLE patients with stable disease (p=0.006) and 18.8±8.0% in SLE patients experiencing a flare (p<0.0001 compared to healthy controls). SAP^+^ TPH levels were significantly greater in SLE patients’ experiencing a flare as compared to SLE patients with stable disease (p=0.04) (Fig. 2C). In SLE, SAP^+^ TPH levels correlated with SLEDAI-2K scores (Fig. 2D) and showed an inverse correlation with complement C4 levels (Fig. 2E).

Given the heterogeneity of SLE disease flares, we explored the association between specific disease activity domains present at visit and the level of SAP^+^ TPH cells in circulation. We found that an expansion of SAP^+^ TPH cells was associated with features of lupus nephritis, such as the presence of nephrotic syndrome and an active urine sediment (defined by >5 RBCs or WBCs per high power field on urine microscopy) on the SLEDAI-2K (Tab. 2).

**Table 2.** Association of SAP^+^TPH levels with SLE SLEDAI-2K Disease Activity Domains.

| SLE Disease Activity Domains: | Association with<br><u>%SAP<sup>+</sup>TPH</u><br><u>p-value</u> |
| --- | --- |
| Arthritis (n=14) | 0.62 |
| Mucocutaneous (n=9) | 0.98 |
| **Lupus Nephritis (n=10):** | **0.05** |
| Proteinuria (>0.5g/day) (n=9) | 0.15 |
| **Urine Sediment (n=4)** | **0.05** |
| **Nephrotic syndrome (n=5)** | **0.02** |
| Alopecia (n=5) | 0.62 |
| Cardiorespiratory: |  |
| Pleural Effusions (n=1) | 0.18 |
| Pericarditis (n=1) | 0.09 |
| Cytopenia (n=4) | 0.90 |
| Low Complements (n=16) | 0.34 |
| **Increased anti-dsDNA (n=19)** | **0.07** |

In a univariable analysis, SAP^+^ TPH levels were significantly greater in the SLE patients with a history of biopsy confirmed LN as compared with SLE patients without renal involvement (19.1±8.0 vs. 13.0±7.2, p=0.04) (Fig. 2F). Importantly, TPH levels without SAP quantification did not differentiate renal from non-renal SLE involvement (23.0±10.2 vs. 16.7±9.0, p=0.10). We analyzed the data using a multivariable model that included age, sex, race, low complement levels and elevated anti-dsDNA antibody levels as potential confounders in the association of SAP^+^ TPH cells with LN. Controlling for these confounders, the level of SAP^+^ TPH cells remained significantly associated with occurrence of LN, such that for every one-point increase in the percent of circulating SAP^+^ TPH cells there was a 3% increase in the odds of having LN (p=0.014) (Tab. 3).

**Table 3.** Multivariable analysis of clinical variables associated with lupus nephritis.

| <u>Lupus Nephritis (LN):</u> | <u>β Coeff</u> | <u>Std Error</u> | <u>p-value</u> |
| --- | --- | --- | --- |
| Age | -0.01 | 0.01 | 0.144 |
| Male Sex | 0.58 | 0.20 | **0.010** |
| Race | -0.01 | 0.12 | 0.936 |
| SAP<sup>+</sup>TPH / CD4 T Cells (%) | 0.03 | 0.01 | **0.014** |
| Low Complement (C3 or C4) | 0.09 | 0.15 | 0.562 |
| Increased anti-dsDNA | -0.03 | 0.23 | 0.882 |

### SAP expression is increased in TFH-like kidney infiltrating T cells in LN

To validate these findings in a different lupus nephritis cohort, we used scRNA-seq data from SLE kidney biopsy samples, available through the AMP collaboration.^17^ Using the original authors’ definitions of the clusters, we compared the expression of the SAP gene, SH2D1A, in T cells from SLE vs. control biopsy samples. SAP expression was higher in SLE T cells as compared to control T cells (p=0.03) (Fig. 3A). We analyzed CD4 and CD8 T cells separately to better understand subset-specific SAP expression patterns. SAP expression across all CD4 T cells did not differ between SLE and control samples. However, SAP expression from healthy kidney CD4 T cells was predominantly seen in T-regulatory cells.

**Figure 3.**
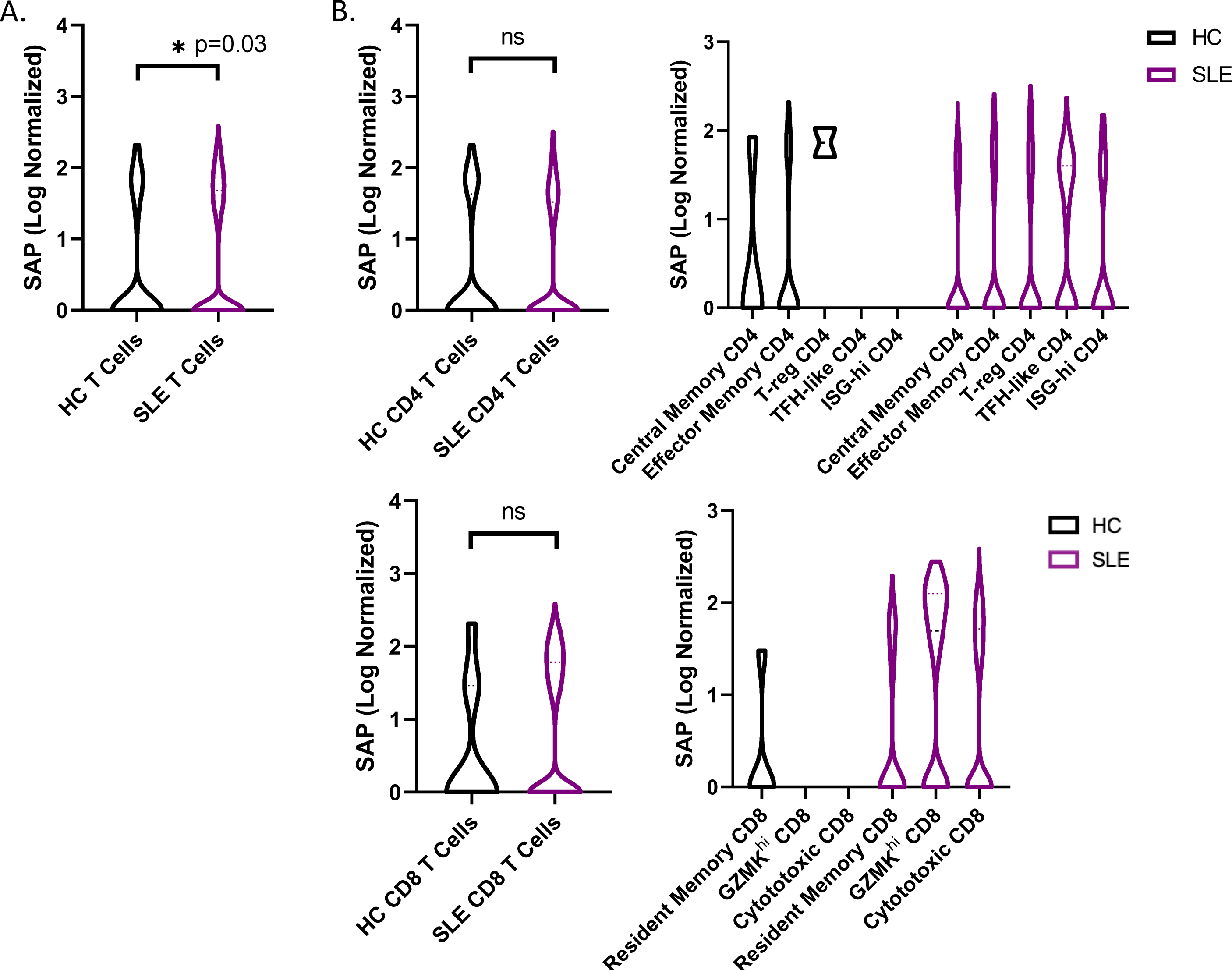
SAP expression in increased in TFH-like kidney infiltrating T cells in LN. Single cell RNA sequencing (scRNA-seq) data from SLE kidney biopsy samples, publicly available through the Accelerated Medicines Partnership (AMP) national collaboration, was analyzed for normalized SAP gene expression. **A)** SAP expression in SLE T cells was compared to that in healthy control (HC) T cells. **B)** Using the original cell cluster annotations as defined by the AMP authors, SAP expression in SLE as compared to HC was stratified by CD4 T cell clusters (top) and CD8 T cell clusters (bottom). T-reg, T regulatory; TFH, T-follicular helper; ISG-hi, interferon stimulated gene-high expression; GZMK, granzyme K. Statistical comparisons were done using a two-sided, unpaired t-test. *p<0.05; ns, non-significant at p=0.05.

In contrast, in SLE kidney T cells, SAP expression was expanded explicitly in the “TFH-like T cell” cluster, the latter annotated based on high expression of CXCR5, PD1, CXCL13, and MAF transcription factor (Fig. 3B top). Similarly, SAP expression across all CD8 T cells did not differ between SLE and control samples. Still, SAP expression in the SLE T cells localized to the cytotoxic and granzyme K (GZMK) expressing CD8 T cell populations that were otherwise not seen in the control kidney T cells (Fig 3B bottom). In summary, SAP expression was increased in kidney infiltrating T cells from LN compared to control kidneys. In LN kidneys, SAP expression was predominantly localized to TFH-like CD4 and GZMK expressing CD8 T cell subsets that were not seen in the control kidney.

We next sought to understand better the transcriptomic signature of LN kidney T cells expressing high as compared to low levels of SAP. We used the normalized SAP gene expression levels to identify the differentially expressed genes between the highest and lowest quartiles of SAP-expressing T cells (Fig. 4A). Increased SAP expression was associated with the expression of SLAMF3 and SLAMF7 and class II antigen-presentation MHC molecules. At the same time, there was differential upregulation of GZMK, GZMA and EOMES, CRTAM, and DTHD1, all of which suggest an activated, cytotoxic CD8 presence. Finally, upregulated CCL4 (MIP-1B), CCL5 (RANTES), CXCR4 and CCR5 cytokine expression were also noted. Interestingly, the IL6R was downregulated in SAP-high T cells. We next analyzed the enriched gene set using KEGG and GO pathway analyses to understand the functions associated with high SAP expression in T-cells better. KEGG analysis identified anti-viral responses, chemokine signaling, cytosolic DNA-sensing cytokine interaction, and Toll-like receptor signaling pathways (Fig. 4B). GO analysis identified enrichment in biological processes associated with T cell activation, MHC Class II peptide presentation, and lymphocyte mediated immunity (Fig. 4C). These results confirmed the presence of SAP expressing “TFH-like” cells in the kidneys of lupus nephritis that was not seen in the control kidney. Additionally, in CD8 T cells, SAP gene expression was seen in the EOMES+ and GZMK+ T cells expressing pro-inflammatory cytokine and chemokine mediators akin to what is commonly seen in anti-viral T-cell activation responses.

**Figure 4.**
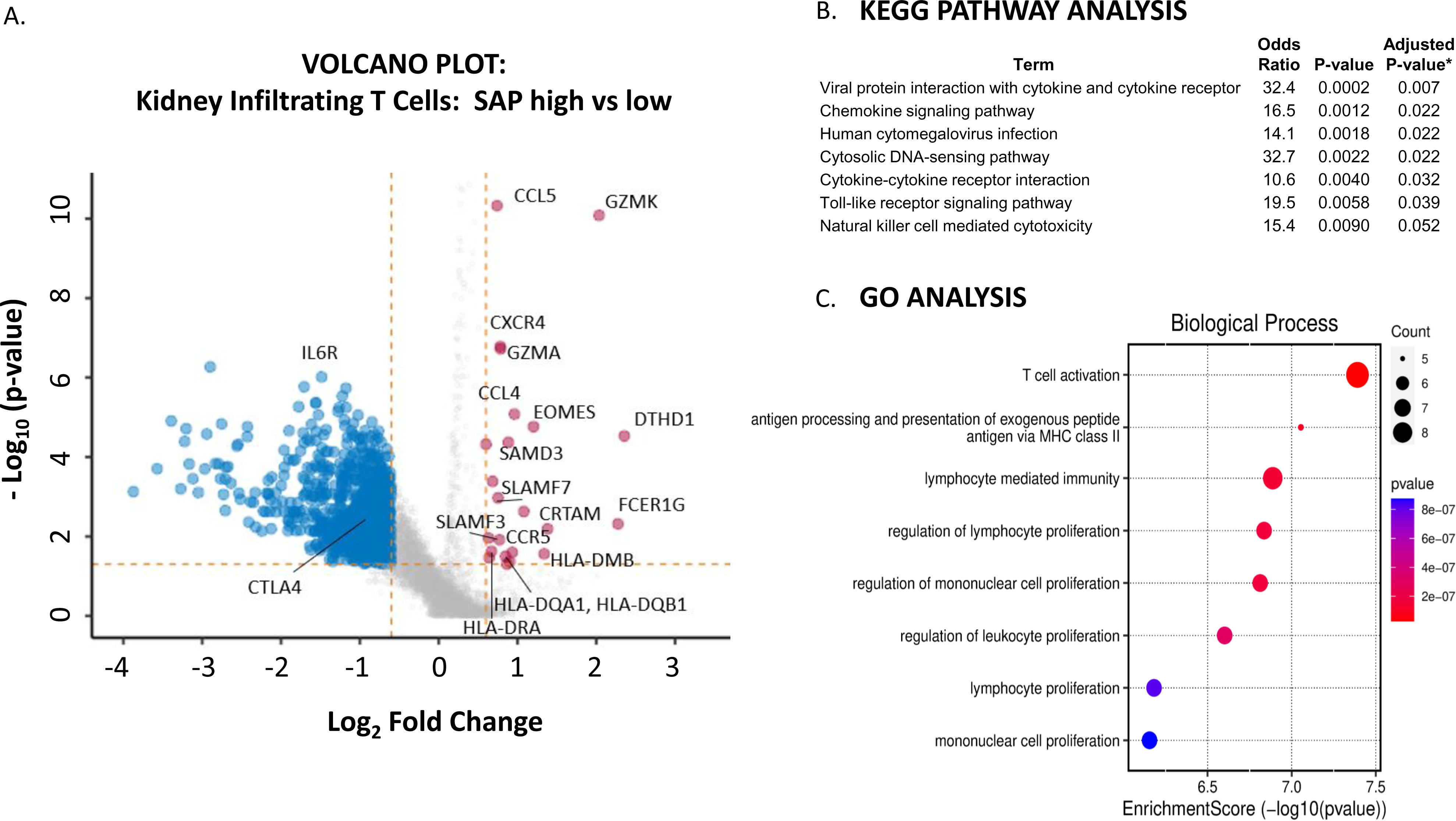
Kidney infiltrating T cells that express high levels of SAP have upregulated expression of genes involved in pro-inflammatory T cell activation. **A)** Differential gene expression of SAP-high as compared with SAP-low kidney infiltrating T cells (n=24 LN and 10 HC samples). The volcano plot shows upregulated and downregulated genes (log2FC ≥ 0.6 and a false discovery rate (FDR) ≥ 0.05). **B)** KEGG Pathway and **C)** GO Analysis of upregulated genes associated with SAP expression. *Adjusted p-value computed using the Benjamini-Hochberg method of correction for multiple comparisons.

#### SAP expression localizes to TFH/TPH-like CD4 T cells and GZMK+ CCL4+ CCL5+ CD8 T cells in an adult human transcriptome of major systemic organs

We next asked whether SAP-expressing T-cell clusters can be isolated from other major human organs. To answer this question, we used single cell T cell RNA-seq data from biopsy samples of fifteen adult organs.^18^ Using this data, we identified 19,495 T cells, which we analyzed using UMAP dimensional reduction and unsupervised Louvain clustering (Fig. 5A). We identified eight clusters of T cells, most of which were non-predominantly distributed amongst the different organs from which they were sampled (Fig. 5B). Restricting the analysis to CD4 T cells, we found that the expression of the SAP gene, SH2D1A, localized predominantly to cluster 8. The additional genes enriched in cluster 8 included ICOS and CD40-ligand and transcription factors MAF and PRDM1, which are consistent with a TFH/TPH-like subset (Fig. 5C).

**Figure 5.**
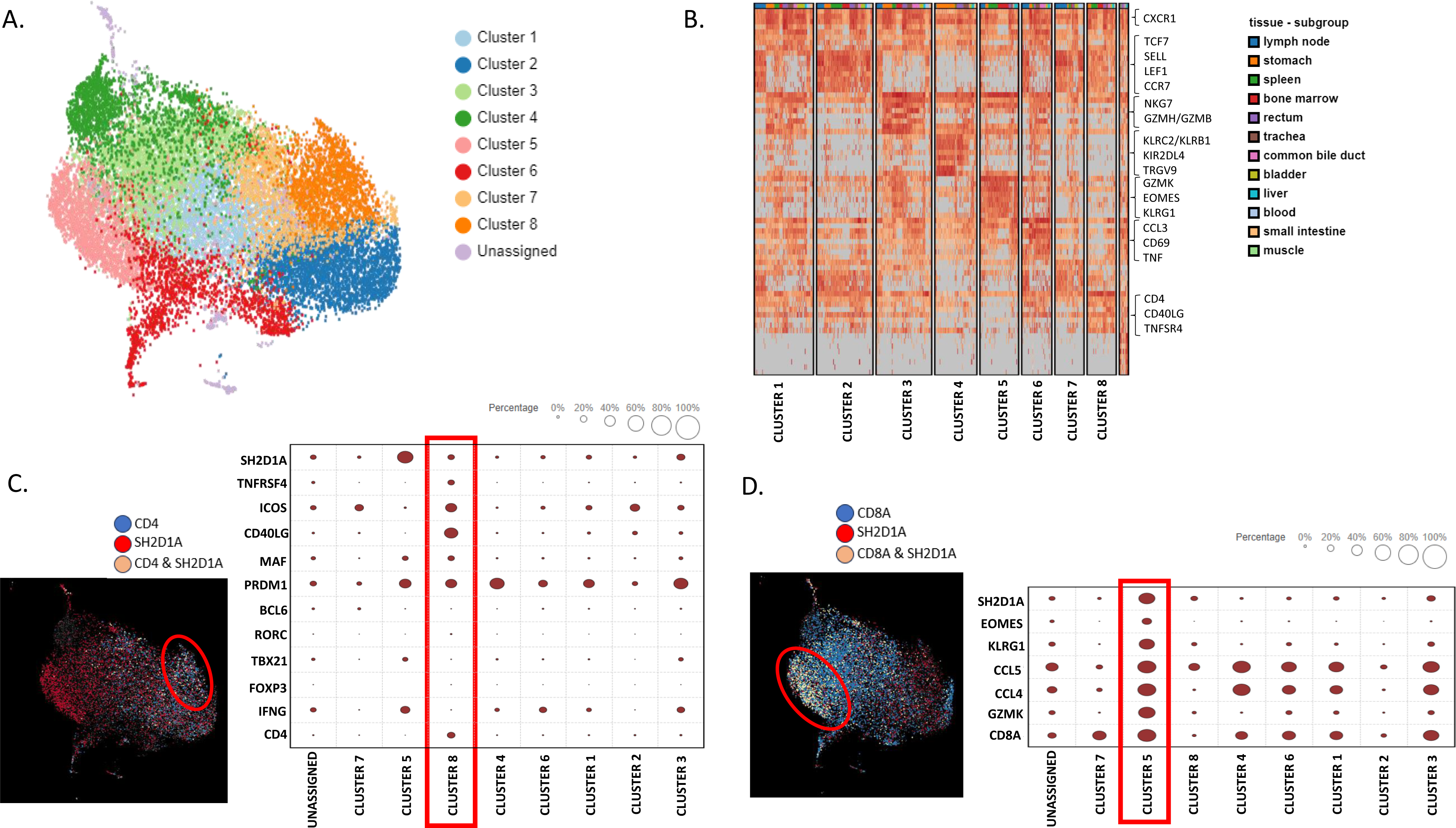
Single cell T cell transcriptome of major human organs confirms two distinct populations of SAP-high T cells, the TFH/TPH-like CD4 and GZMK+EOMES+ effector CD8. **A)** Dimension reduction of 19,495 T cells was performed using the Uniform Manifold Approximation and Projection (UMAP) algorithm, followed by Louvain clustering and **B)** heat map of major cell subsets. Columns represent selected differentially expressed signature genes in each cluster, and different clusters are exhibited in the rows. **C)** Cellular composition of SAP gene (SH2D1A) and CD4 co-expression (left) mapped to cluster 8. Selected differentially expressed signature genes quantified (right). **D)** Cellular composition of SH2D1A and CD8 co-expression (left) mapped to cluster 5. Selected differentially expressed signature genes quantified (right).

Conversely, restricting the analysis to CD8 T cells, we found SAP expression in cluster 5. Enriched genes from this cluster included EOMES, GZMK, KLRG1, CCL4, and CCL5 (Fig. 5D). We were therefore able to confirm that organ infiltrating T cells, even in absence of SLE, have two distinct populations of SAP-high T cells: the TFH/TPH-like CD4 and EOMES+, GZMK + CD8 T subsets. These organ-infiltrating T cells isolated from a healthy volunteer suggest a baseline T cell presence that has the potential to become expanded in the context of SLE disease.

## Discussion

SLAMF receptors are important co-stimulatory TCR receptors critical for T-cell mediated B-cell development. In this work we found elevated SAP levels in T cells from SLE patients compared to healthy controls. SAP levels correlated with markers of T cell activation, such as CD69 and HLA-DR expression. The highest levels of SAP expression in CD4 T cells were seen in the TPH, followed by the TFH, subsets.

TPH was first described in the synovium of patients with rheumatoid arthritis and has since been found to be expanded in systemic lupus, where its numbers correlate with disease activity and autoantibody titers.^12, 13, 20^ These cells were also identified in the kidneys of patients with lupus nephritis, where they correlated with pathogenic B cell subsets.^21^ In our SLE cohort we found that SAP^+^TPH cells, but not total TPH cells, were associated with biopsy-proven LN. We believe that SAP^+^TPH cells identify a subset of functionally active TPH cells poised to support B cell maturation in the periphery more readily. SAP^+^TPH cells were also associated with active SLE (higher disease activity scores measured by SLEDAI-2K) and showed a trend towards an inverse correlation with complement C3 and C4 levels. The association between SAP^+^TPH cells and LN remained significant in multivariable analyses, controlling for common confounders such as age, sex, race, anti-dsDNA antibody levels, and complement levels. Additionally, using scRNA-seq data from kidney biopsy samples of patients with LN, we validated the presence of TPH-like T cells that express high levels of SAP mRNA and are further expanded in SLE compared to control.

The SAP gene (SH2D1A) is expressed on the X chromosome in humans. Loss of function SAP mutations result in disorganized germinal centers and lack of antigen-specific B cell differentiation, clinically manifesting as an X-linked immunodeficiency syndrome. Conversely, we propose that increased SLAM/SAP signaling functions augment T-B cell interactions and enhance autoantibody formation – conferring a risk for SLE. A single nucleotide polymorphism in the SLAMF3 gene, which changes the conformation of the SLAMF3 receptor to have a more vital molecular interaction with SAP, is associated with the development of SLE.^3, 22^ Mutations in the SAP gene associated with SLE risk have also been reported, although the functional significance of these mutations is not well understood.^23^ On the other hand, the discovery of a de-novo frameshift mutation in the SAP gene of SLE-prone mice ameliorated the predisposition of these mice to develop SLE.^11^ Similarly, repression of SAP gene transcription by a transcription factor myocyte enhancer factor 2 (Mef2d) has been shown to inhibit SAP-dependent T-B synapse formation and prevent antigen-specific CD4 T cells from differentiation into germinal center TFH cells. MEF2D mRNA expression inversely correlates with TFH populations, autoantibodies, and SLEDAI scores in SLE.^24^ Our finding that SAP levels are increased in the circulation of SLE T cells, predominantly TFH and TPH cells, further supports the immunopathogenic role of SLAM-SAP signaling in SLE.

Our findings of increased SAP levels in SLE contrast with the work by Karampetsou et al. reporting decreased SAP in SLE T cells as compared to controls.^25^ There are two possible explanations for this. First, we suspect that there is clinical and molecular heterogeneity in SLE but organ specific SLE involvement is not considered by Karampetsou et al. Second, SAP levels in T cells are rapidly downregulated following T cell activation, thus difference in the processing/activation of PBMCs can lead to markedly different conclusions.^26^ Indeed, when Karampetsou et al. analyzed T cell lysates at different time-points, the decrease in T cell SAP levels in SLE was only seen after 5 hours of T cell culture treatment. Given that we found a correlation between SAP levels and markers of T cell activation (CD69, HLA-DR) it is possible that SAP level degradation over time in culture can account for the different observations in our study vs. Karampetsou et al.

Growing evidence suggests that tubulointerstitial inflammation is a strong predictor of long-term outcomes in lupus nephritis.^27, 28^ T cells compose more than 60% of all immune cells in the tubulointerstitium, and aggregates of T and B cells are seen in up to 50% of biopsy specimens.^29, 30^ It is believed that peripheral T and B cells traffic to the renal interstitium, where they become organized into tertiary lymphoid-like structures, promoting B cell isotype switching and in-situ auto-antigen responses. The expanded SAP^+^TPH cells in the circulation of patients with LN and the presence of these TPH-like cells in the kidney biopsy samples suggest a plausible pathological trajectory for these cells from the periphery to the lupus kidney. Furthermore, a possible target B cell population of TPH includes the CD11c^+^CD21^-^ CXCR5^-^ age-associated B cells (ABC). This pro-inflammatory B cell subset rapidly expands and differentiates into plasmablasts upon activation.^31^ Indeed, the CD11c^+^ B cells in circulation correlate with markers of SLE disease activity and specifically with the presence of lupus nephritis; the same cells are also found in the tissues of nephritic kidneys where they localize to ectopic lymphoid follicles.^32, 33^ It is thus exciting to consider that the SAP^+^TPH subset identified by us to be associated with LN may represent a cognate partner to the ABC – with the possibility that both cell types can migrate into the target tissue, organize into ectopic germinal cells, and contribute to the development of LN.

The differential gene analysis of SAP-high vs. SAP-low-expressing kidney infiltrating T cells confirmed that SAP expression was associated with members of the SLAMF family, specifically SLAMF3 and SLAMF7. However, while the role of SAP is well described in the literature as being critical for CD4 T-B cell-mediated interactions, we additionally found that SAP expression was also strongly associated with a pro-inflammatory, cytotoxic CD8 effector T cell phenotype based on the high expression of GZMK, EOMES, DTHD1, and CCL4 (MIP-1) and CCL5 (RANTES). We found that the TPH-like CD4 subset and the GZMK+EOMES+ CD8 subset, expressing high levels of SAP, can be isolated from many healthy adult organs. Still, these cell populations were expanded in the kidneys of patients with lupus nephritis compared to control kidneys. The presence and function of the organ-infiltrating SAP-positive CD8 T cells in the steady state, and further expanded in inflammatory disease, still needs to be better understood. Given the gene enrichment of MHC-Class II antigen presentation genes in SAP-expressing T cells, it is possible that SLAM-SLAM signaling is not only critical for CD4 T-B cell signaling but also for in-situ CD4 T-cell assisted maturation of cognate CD8 T cells. Additionally, while there are multiple SLAM family receptors that share functional redundancy, the downstream signaling in the T cell converges on binding SAP. SAP then functions to recruit kinases that enhance phosphorylation of downstream TCR signaling proteins (i.e., ZAP-70, LAT) and thus T cell activation.^8^ Accelerated phosphorylation downstream of TCR enhances T cell hyper-responsiveness to self-ligands, suggesting increased SAP levels may have a mechanistic role in lowering the threshold of T-cell reactivity to self-antigen, thus initiating auto-reactive adaptive immunity.^34^ In this case, pharmacological disruption of the SAP signaling pathway may specifically target the more pathogenic helper and effector lymphocyte subsets in SLE (Fig. 6).

**Figure 6.**
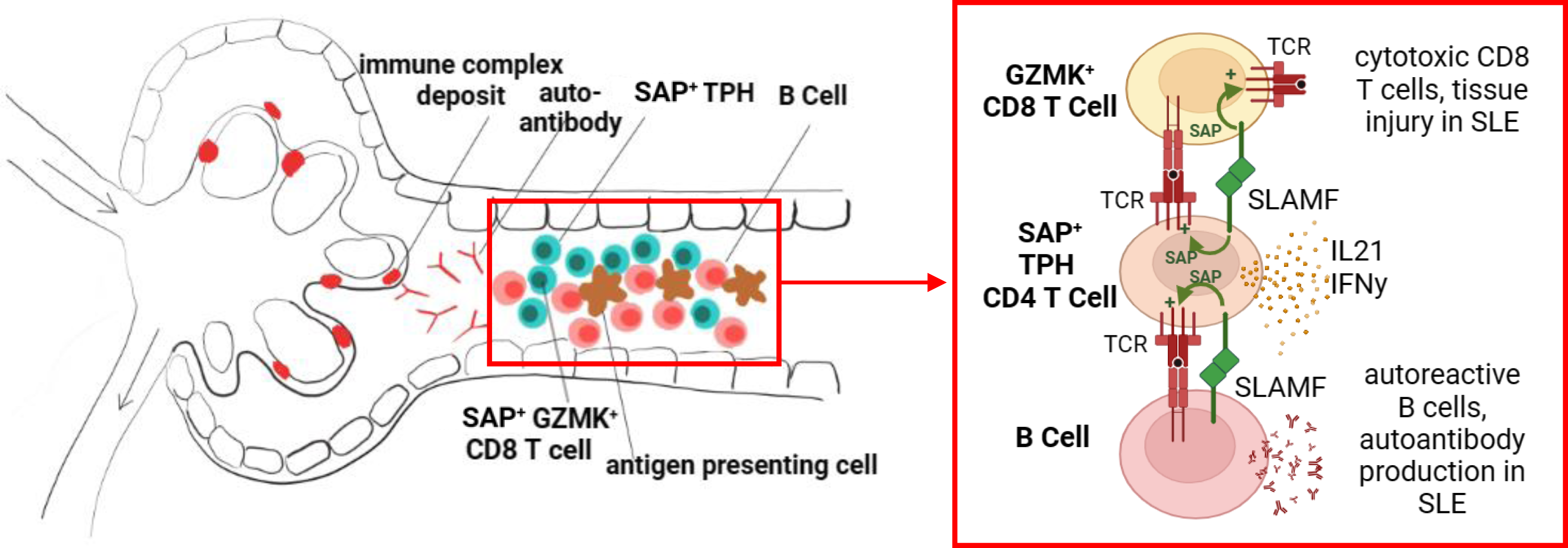
SLAM/SAP signaling is hypothesized to contributes to formation of tertiary lymphoid structures and in-situ development of autoreactive lymphocytes. **A.** We propose that the expanded SAP-positive TPH cell population in SLE traffics to inflamed tissues, such as the kidney tubulointerstitium, and promotes B cell and cytotoxic CD8 T cell development in situ. **B.** TPH cells support developing CD8 and B cells via direct cell-cell interactions, dependent on SLAMF-SLAMF binding and downstream SAP signaling, as well as release of cytokines such as interferon gamma (IFNy) and interleukin 21 (IL-21). The images were created with BioRender.com.

Our work has several limitations. We studied a small number of patients and future work will need to validate these findings in a larger sample. Additionally, as SAP is not expressed by B cells, we focused our analyses on T cells. As a result, we cannot make any conclusions about direct B cell activation.

Finally, we used the lupus kidney scRNA-seq data available from the AMP consortium but given the relatively small number of T cells isolated from the kidney biopsy samples, we could not separately analyze CD4 from CD8 T cell clusters. Instead, we had to broaden the differential gene analyses to all T cells. As a result of this, compounded by the overall low-depth sequencing of the SAP gene, we tolerated a lower log2FC cutoff to increase sensitivity in discovering novel signaling molecules associated with high SAP expression. The analysis of the KEGG pathway in this context links the identified SAP-associated genes with Toll-like receptor signaling and cytosolic DNA-sensing responses, both of which are strongly linked to monogenic as well as polygenic SLE risk.

In summary, we have identified an expanded SAP^+^TPH population in the circulation and kidney samples of patients with biopsy-confirmed lupus nephritis. We believe this is a functionally active TPH cell population directly involved in in situ ectopic germinal center formations. We also identified a GZMK^+^ SAP^+^ cytotoxic CD8 T cell population with a pro-inflammatory cytokine and chemokine gene expression. The role of these T cell subsets in promoting direct tissue damage in lupus nephritis will be the subject of future research.

## Author Contributions

The project was designed and supervised by YG, ADA and AM. Clinical samples were collected by YG, LGP, and LK. YG, SB, SL, ADA and AM contributed to data analysis. The paper was written by YG. All authors contributed to the article and approved the submitted version.

## Acknowledgments

None

---

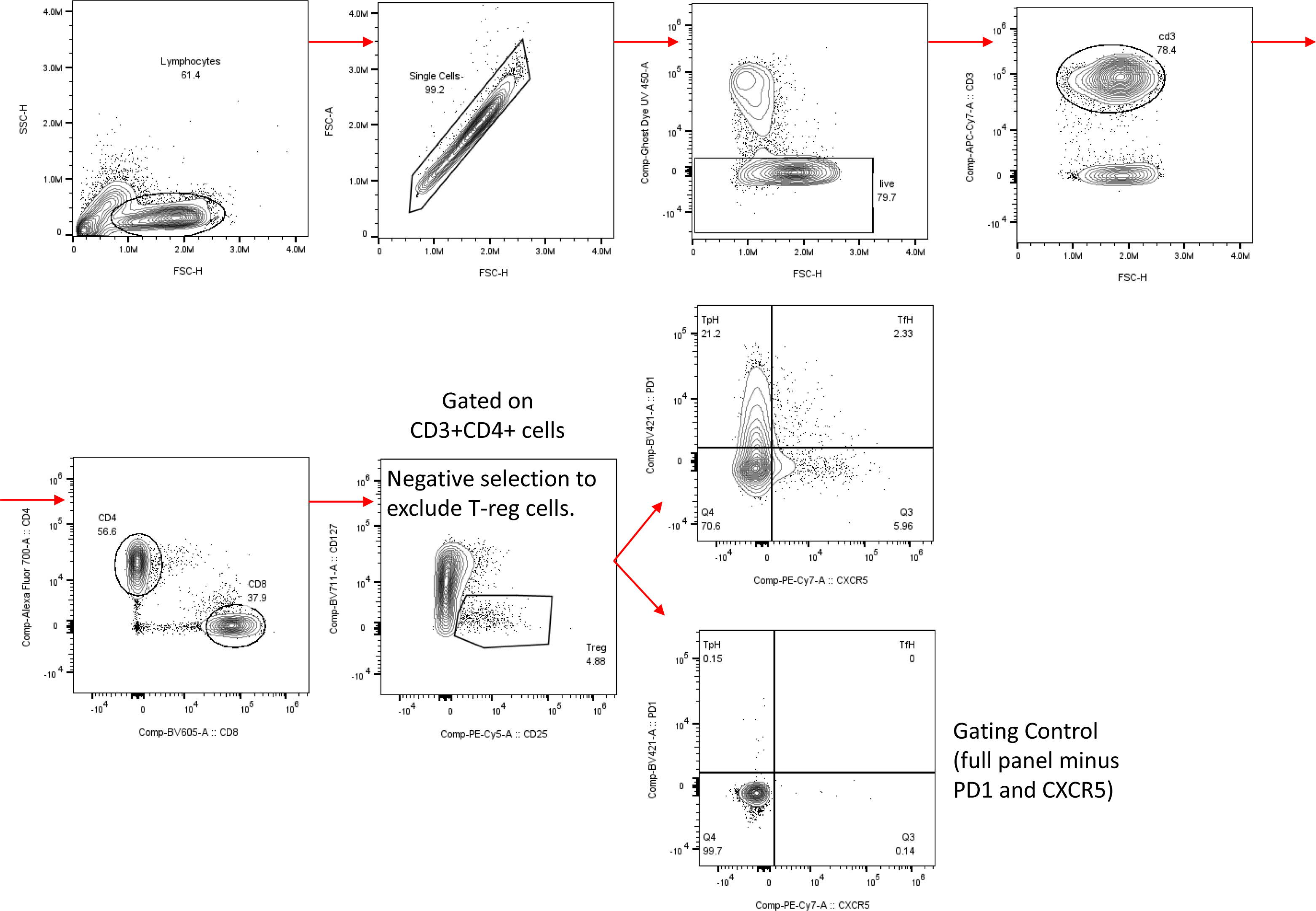

